# A Novel Neoantigen Discovery Approach based on Chromatin High Order Conformation

**DOI:** 10.1101/539387

**Authors:** Yi Shi, Mingxuan Zhang, Luming Meng, Xianbin Su, Xueying Shang, Qingjiao Li, Mengna Lin, Xin Zou, Qing Luo, Yangyang Zhai, Kunyan He, Lan Wang, Cong Chen, Xiaofang Cui, Na Wang, Jian He, Yaoliang Yu, Yanting Wu, Lintai Da, Tom Weidong Cai, Ze-guang Han

## Abstract

High-throughput sequencing technology has yielded reliable and ultra-fast sequencing for DNA and RNA. For tumor cells of cancer patients, when combining the results of DNA and RNA sequencing, one can identify potential neoantigens that stimulate the immune response of the T cell. However, when the somatic mutations are abundant, it is computationally challenging to efficiently prioritize the identified neoantigen candidates according to their ability of activating the T cell immuno-response. Numerous prioritization or prediction approaches have been proposed to address this issue but none of them considers the original DNA loci of the neoantigens from the perspective of 3D genome. Here we retrospect the DNA origins of the immune-positive and non-negative neoantigens in the context of 3D genome and discovered that 1) DNA loci of the immuno-positive neoantigens tend to cluster genome-wise; 2) DNA loci of the immuno-positive neoantigens tend to belong to active chromosomal compartment (compartment A) in some chromosomes; 3). DNA loci of the immuno-positive neoantigens tend to locate at specific regions in the 3D genome. We believe that the 3D genome information will help to increase the precision of neoantigen prioritization and discovery and eventually benefit precision and personalized medicine in cancer immunotherapy.

## 1 Introduction

In a variety of human malignancies, immunotherapies via boosting the endogenous T cell’s ability to destroying cancer cells have demonstrated therapeutic efficacy[1]. Based on clinical practices in a substantial fraction of patients, the inference of endogenous T cell with mounted cancer-killing ability is that the T cell receptor (TCR) is able to recognize peptide epitopes that are displayed on major histocompatibility complexes (MHCs) on the surface of the tumor cells. These cancer rejection epitopes may be derived from two origins: the first origin of potential cancer rejection antigens is formed by non-mutated proteins to which T cell tolerance is incomplete for instance, because of their restricted tissue expression pattern; the second origin of potential cancer rejection antigens is formed by peptides that cannot be found from the normal human genome, so-called neoantigens[1]. With the development of genome sequencing, it has been revealed that during cancer initiation and progression, tens to thousands of different somatic mutations are generated. Most of these mutations are passenger mutations, meaning there is no obvious growth advantage, and are often caused by genomic instability within the tumor cells. A limited number of cancer mutations are driver mutations which interfere with normal cell regulation and help to drive cancer growth and resistance to targeted therapies[2]. Both passenger mutations and driver mutations can be nonsynonymous that alter protein coding sequences, causing tumor to express abnormal proteins that cannot be found in normal cells. When cell metabolize, the proteins possessing abnormal sequences are cut into short peptides, namely epitopes, and are presented on the cell surface by the major histocompatibility complex (MHC, or human leukocyte antigen (HLA) in humans) which have a chance to be recognizable by T cell as foreign antigens[2].

According to the discoveries mentioned above, in theory therefore, if the potential neoantigens can be identified via sequencing technology, one can synthesize epitope peptides in vitro and validate their efficacy in vivo (cancer cell-line or in mouse model) before clinical practice[1, 2]. Indeed, cancers with a single dominant mutation can often be treated effectively by targeting the dominant driver mutation[2, 3]. However, when the somatic mutations are abundant, which is the case in most cancer types, it is computationally challenging to efficiently prioritize the identified neoantigen candidates according to their ability to activate the T cell’s immuno-response[4]. Over the past few decades, numerous neoantigen prediction approaches have been proposed to address this issue[5-7]. These approaches can be classified into two major categories: the protein 3D structure-based approaches which consider the pMHC and TCR 3D conformation, and the protein sequence-based approaches which consider the amino acid sequence of protein antigens. For the protein 3D structure-based approaches, in some specific cases when high quality pMHC 3D structures are available, molecular dynamic (MD) methods are used to explore the contact affinity of pMHC-TCR complex[8-10], in most cases, however, the modelling or simulation by protein docking and threading has to be used due to the lack of high quality pMHC 3D structures. Most approaches belong to the sequence-based category as there are much larger data sets for training and validation[11, 12] and because they are usually very efficient to set up[4, 13].

Early sequence-based methods relied on position-specific scoring matrices (PSSMs), such as BIMAS[14] and SYFPEITHI[15], in which the PSSMs are defined from experimentally confirmed peptide binders of a particular MHC allele[4]. Later, more advanced methods based on machine-learning techniques have been developed to capture and utilize the nonlinear nature of the pMHC-TCR interaction which indeed demonstrated better performance than the PSSM-based methods. Consensus methods that combine multiple tools to obtain more reliable predictions were also developed, such as CONSENSUS[16] and NetMHCcons[17], which demonstrated better performances; for these methods however, the performance gain is determined by the weighting scheme which cost increased computational power. When considering peptide binding, most methods did not consider the HLA allele variety, therefore, pan-specific methods, such as NetMHCpan[6, 7], are developed which allow the HLA type independent prioritization.

As one of the widely adopted practices in neoantigen prioritization, NetMHCpan first trains a neural network based on multiple public datasets, then the affinity of a given peptide-MHC considering the polymorphic HLA types HLA-A, HLA-B or HLA-C is computed according to the trained neural network. NetMHCpan[7] and NetMHCIIpan[18] perform remarkably, even compared to allele-specific approaches[4, 19]. However, although several assessments and criteria were proposed in the past aiming at a more fair and effective comparison[19-21], there are no recent independent benchmark studies that can be used to recommend specific tools up until now. More importantly, however, to the best of our knowledge, none of the neoantigen prediction methods mentioned above consider the mutation DNA loci of the neoantigens in the perspective of 3D genome, which carries much richer information compared to the amino acid sequence alone[22]. In this work, we retrospect the DNA origin of the immune-positive and non-negative neoantigens in the context of the 3D genome and demonstrate some discoveries that worth paying attention to.

## 2 Methods

### 2.1 Data collection and curation

All the peptide sequences and their corresponding immune effectiveness were collected from IEDB (T Cell Assay)[12] on May 27^th^ 2018; the raw dataset contains 337248 peptide records. We narrowed down to Homo sapiens and MHC-I subtypes and then further restrain the AA length to be equals to 9 with duplicated peptide merged. Finally, we obtained 3909 qualified records with 809 immuno-positive peptides and 3100 immuno-negative peptides that has mapping hits in the human hg19 reference genome. Note that for identical peptides with multiple immune experiments, we define positive rate > 0.8 as immune-positive peptides and positive rate < 0.2 as immune-negative peptides. In detail, there are two steps in this procedure:

Step I. Extracting the Homo sapiens peptide sequences and cleaning up the dataset from initial dataset. We used PANDAS library to create a data frame object for processing. Then we assigned the column name by importing a name dictionary. Then we filtered the dataset so that the only entries left have “homo sapiens” as their hostname, after which we transfer the dataset to a tsv file for the future procedures. Then, we further clean up the dataset by introducing two helper functions: 1. Letter_check takes in a string as the parameter and checks for letters that are not in the alphabet for a proper peptide sequence. The function returns an indicator which value is set to true if the string is a legal peptide sequence. 2. Drop_legal takes a data frame, and iterate through and deletes the rows with illegal strings. With these helper functions, we can remove all the illegal entries in the dataset.

Step II: Counting the number of sequences that have positive qualitative measure. The algorithm here takes advantage of the layout of the dataset. We keep counting until the last appearance of a certain sequence; add 1 to the counter of positivity if a positive qualitative measure is detected. For the last appearance of the sequence, we can either add 1 if its positive or we skip to the next step. The counter resets every time we finish counting positivity for a sequence and move on to the next one. We store the counted values into a dictionary that have sequence and MHC as its key.

For the chromatin 3D conformation data, we used the Hi-C data of hESC and IMR90 cell lines generated by Bin Ren’s lab[23]. The contact frequencies and the subsequent chromatin 3D modeling are based on these Hi-C data.

### 2.2 Mapping peptides to human genome

To map the peptides to human genome (hg19), we wrote a pipeline to query the BLAST[24] web server and map the gene names to chromosomes and starting positions. The algorithm first divided the dataset into 711 folds where each fold has 100 sequences for the BLAST server to process. To set up the BLAST search, we regulated the searching algorithm to search for Homo sapiens matches only with entrez ID keywords and used the PAM30 matrix to search for matches. We also adjusted the gap costs to regulate gap penalty. After the setup, we called BLAST iteratively and wrote the result onto a tsv file. For each match, we saved the accession and raw bit score for the first hit. After acquiring the accessions, we uploaded a list of refseq id to the DAVID tool[25] to obtain the gene names composed with gene symbols. The algorithm mapped gene names to chromosome positions, and we started with a dataset that records chromosome positions and gene names for numerous genes as our database. To save time during iterations, we created two dictionaries recording chromosome positions with gene names as keys, one from the dataset we produced from BLAST results and one from the database. We iterated through the dictionaries simultaneously. If we found a match for the keys, we recorded the chromosome positions on the result file. The final result is in the form of a tuple that contains peptides, HLA subtype, chromosome number, and chromosome position.

### 2.3 Active and inactive compartment determination

To determine the compartment activeness (compartment A: active, compartment B: inactive) of each chromosome bin, we used individual chromosome Hi-C contact maps. We first diagonal normalized each contact map by dividing the contact frequencies by their corresponding off-diagonal mean. Then we computed the Pearson correlation coefficient (PCC) matrices for each chromosome, and the compartment type was jointly determined by the sign of eigenvalue of the first eigenvector of the PCC matrices and the signal of the epigenetic marker H3k4me1.

### 2.4 Chromatin 3D modeling

We developed a new method for modeling 3D conformations of human genome using molecular dynamics (MD) based approach with resolution of 500kb (bin size) for hESC and IMR90 Hi-C data. In this method, each bin was coarse-grained as one bead and intact genome was modeled as 23 polymer chains represented by bead-on-the-string structures. The spatial position of each bead is affected by two factors: (1) chromatin connectivity that constrains sequentially neighbor beads in close spatial proximity and (2) chromatin activity that ensures active regions are more likely to be located close to the center of cell nucleus. In this work, chromatin activity was estimated as compartment degree that can be directly calculated from Hi-C matrix with algorithm described in previous work[26]. Based on the relative values of compartment degrees, all the beads were assigned distances with different values to nuclear center and then the conformation of chromatin was optimized from random structures with molecular dynamics approach by applying bias potential to satisfy these distance constraints. For each cell linage, 300 conformation replicas were optimized from random ones to reduce possible bias for further analysis.

## 3 Results

### 3.1 Neoantigen proximity in individual chromosome (intra-chromosome)

We generated all peptide pairs between immune-positive peptides and peptide pairs between immune-negative peptides. Then on each chromosome (intra-chromosome), we generate each pair’s contact frequency on hESC and IMR90 Hi-C data[23]. The results are shown in Figure 1a and Figure 1b. Jointly from these results, we found that positive peptides’ corresponding DNA loci tend to be more proximate (p<0.05) than the negative ones on chromosome 1 (chr1), chr7, chr10, and chr12, while negative peptides’ corresponding DNA loci tend to be more proximate than the positive ones on chromosome chr2, chr5, chr8, chr11, and chr20.

**Figure 1.**
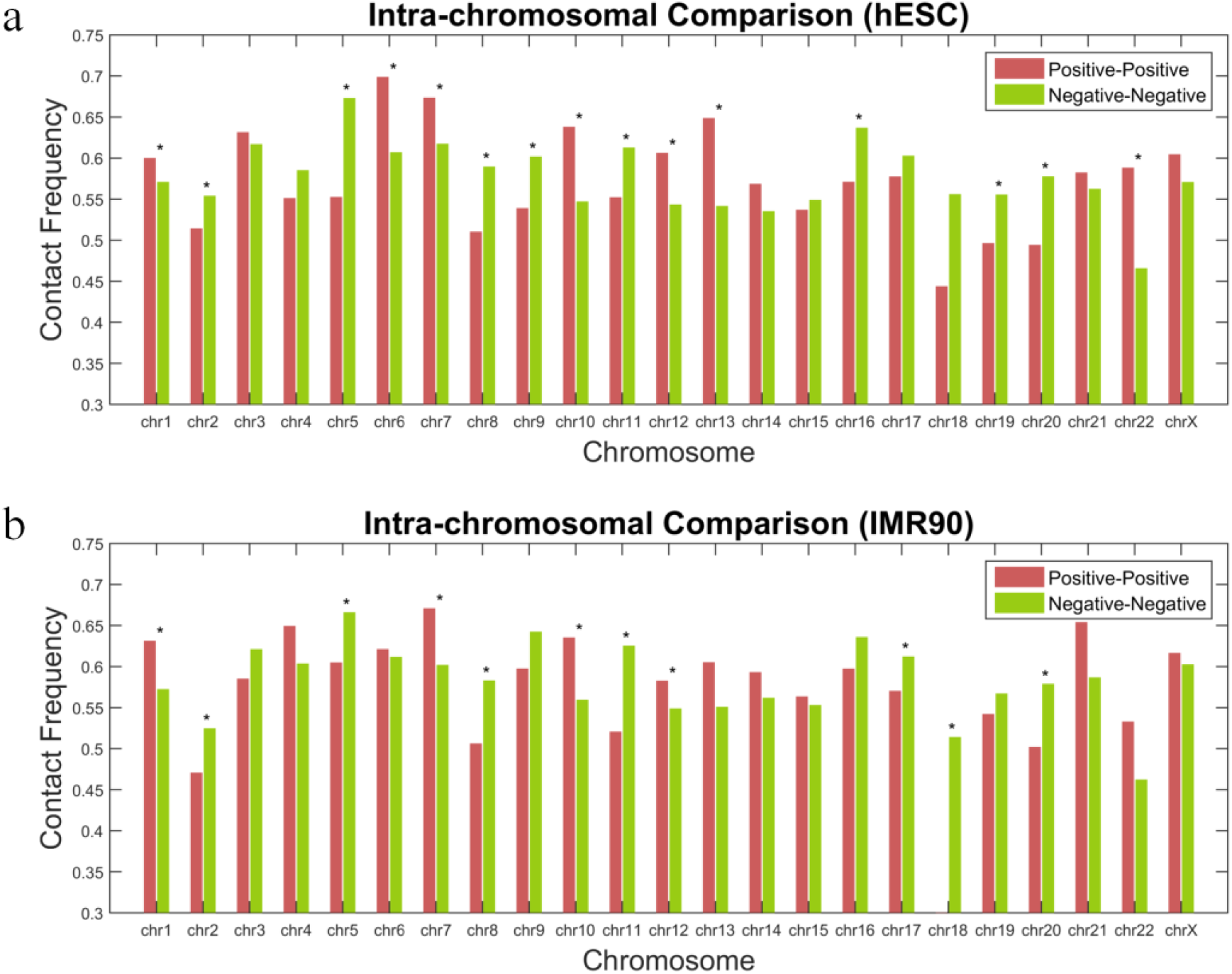
Average contact frequency (CF) of immune-positive peptide pairs and immune-negative peptide pairs based on (a) hESC Hi-C data and (b) IMR90 Hi-C data. The star sign indicating p<0.05.

### 3.2 Neoantigen proximity in the whole genome (inter-chromosome)

For the inter-chromosomal peptide pairs, both positive and negative, we also collect their contact frequency and calculate the average values. As shown in Figure 2, on both hESC and IMR90 Hi-C data, immune-positive peptide pairs are more proximate to each other comparing to immune-negative peptide pairs. The corresponding P-values are close to zero.

**Figure 2.**
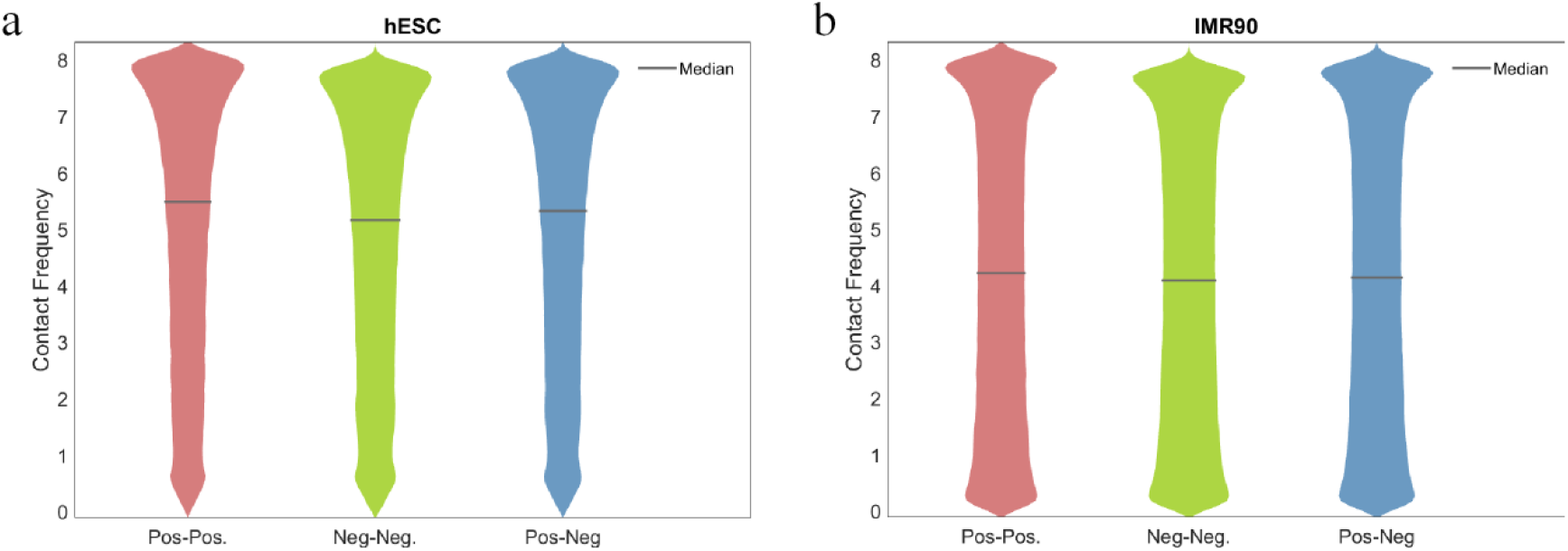
Contact frequency distribution comparison between immune-positive peptide pair’s DNA loci, immune-negative peptide pairs’ DNA loci, and immune-positive-negative peptide pairs’ DNA loci, on (a) hESC Hi-C data and (b) IM4R90 Hi-C data.

### 3.3 Neoantigen distribution on active and inactive compartment

For each chromosome, we compute the compartment type A/B (A: active; B: inactive) for each chromosomal region (bin), shown in Figure 3a and 3b. Then we assign positive and negative peptides with their corresponding A/B compartment type. We found that in certain chromosomes, immune-positive neoantigens tend to be located on compartment A, comparing to immune-negative neoantigens, as shown in Figure 3c.

**Figure 3.**
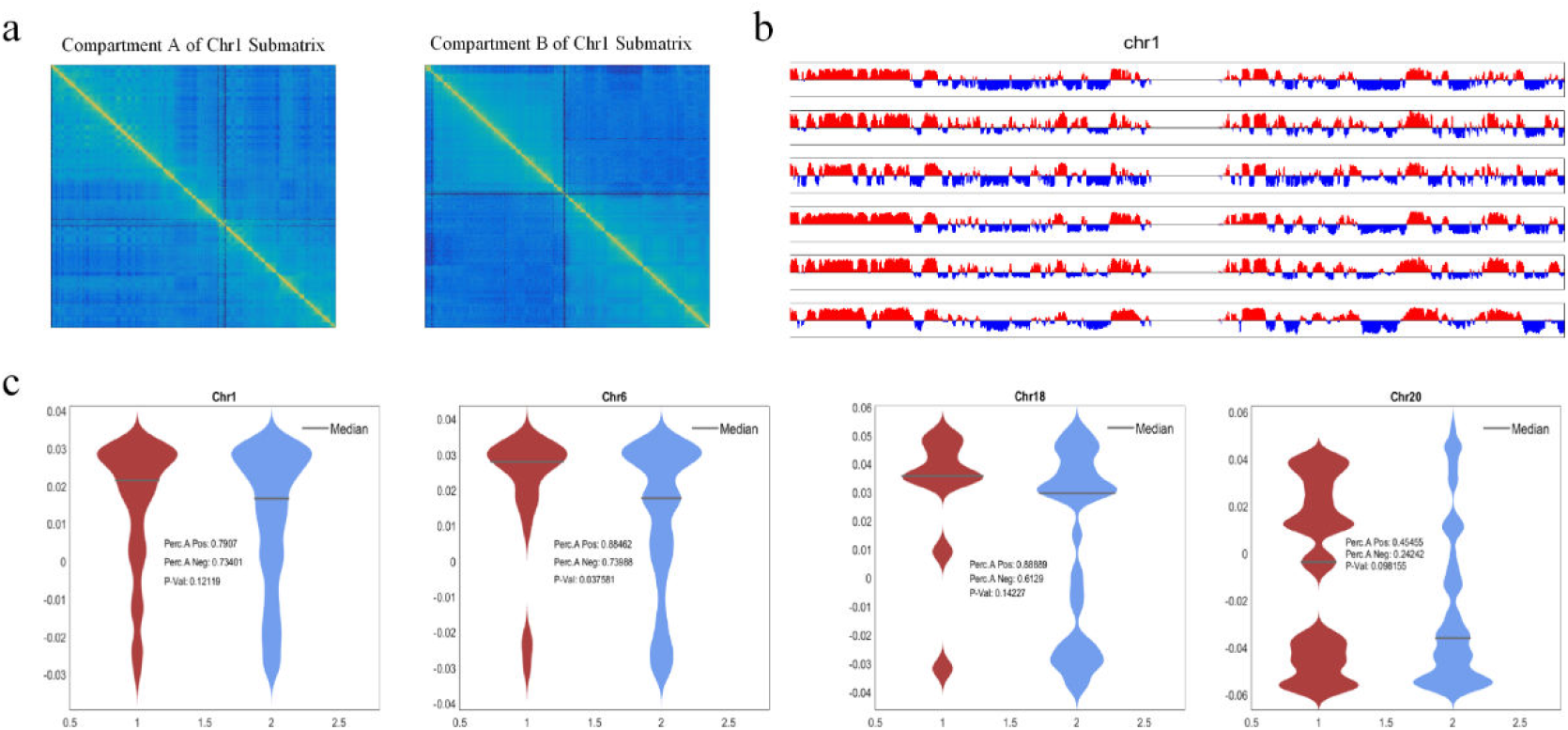
Neoantigen distribution in active and inactive compartments. (a) Example of compartment A and compartment B submatrix on chromosome 1. (b) Example of compartment A/B values on chromosome 1. (c) Distribution of percentage of compartment A of immune-positive (red) and immune-negative peptides (blue).

### 3.4 The radius position distribution of neoantigen

We developed a novel molecular dynamic based approach to model the 3D conformation of the human genome, on both hESC and IMR90 Hi-C data. We then mapped the positive and negative peptides’ corresponding chromosomal loci on the constructed 3D genome and calculated their radius distance to the nucleus center, as shown in Figure 4a. We found that the immune-positive peptide’s corresponding loci tend to locate closer to the nuclear periphery, compared to the negative ones, as Figure 4b demonstrates. We then used the radius position as the immunogenicity predictor and found that combined with existing widely adopted methods such as netMHCPan and netMHC, the joint prediction, defined as *Y_pred_* = *S_netMHCPan_* × *r*^2^ or *Y_pre_* × *S_netMHC_* × *r*^2^ can discriminate the immune-positive peptides from the immune-negative peptides. As shown in Figure 4c, the AUPR increased from 0.55 to 0.64. The AUC of the ROC curves are also increased.

**Figure 4.**
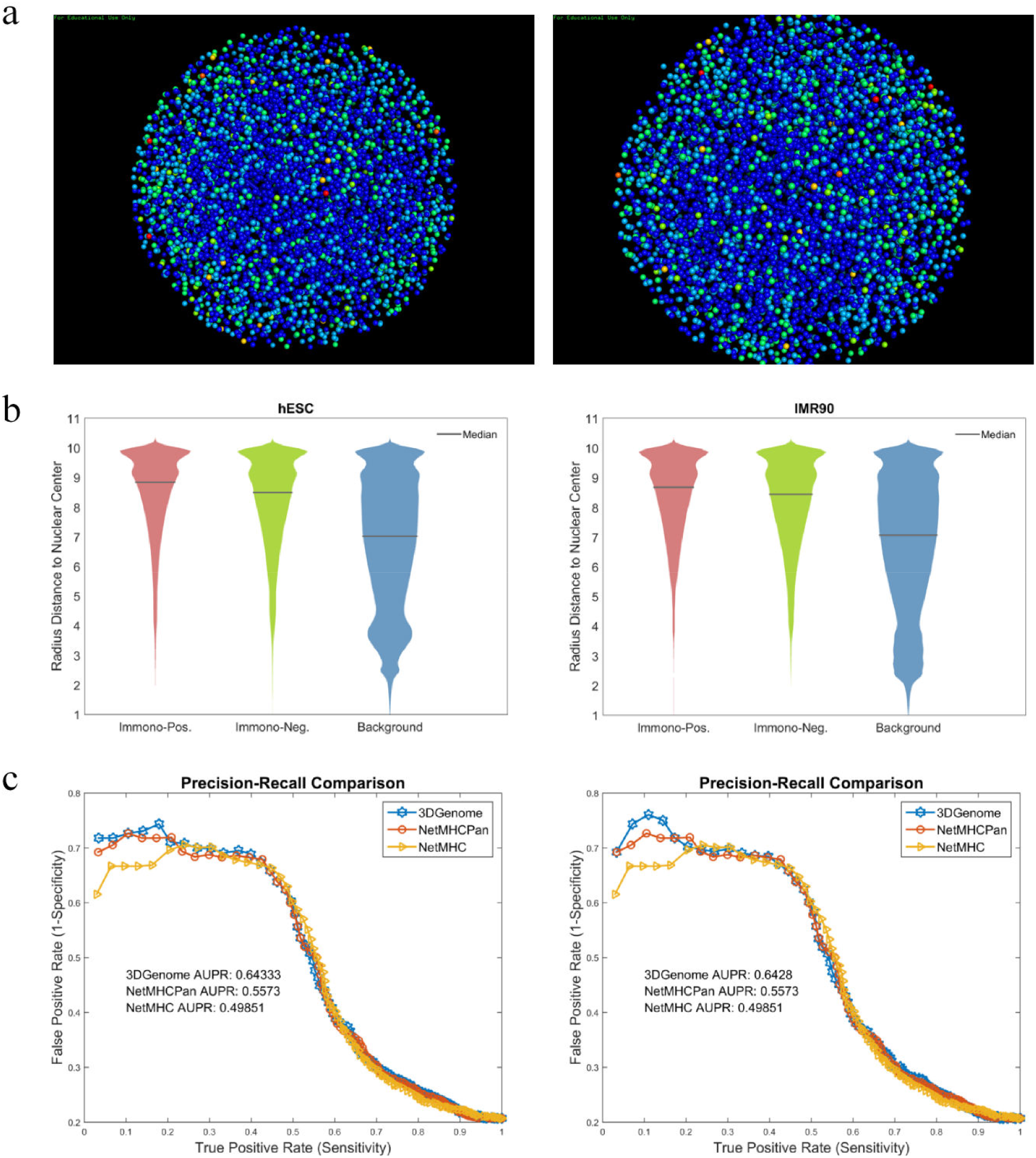
The radius position distribution of neoantigen. (a) Positive and negative peptides’ corresponding chromosomal loci on 3D genome structure. (b) Radius position distribution of the positive peptides comparing to the negative peptides. (c) Precision-recall curve demonstrating the discriminative power of radius position in immunogenicity prediction.

## 4 Discussion

In cancer immune therapy, neoantigen therapy is a rising and promising topic as it can be genuinely personalized and precise. However, when the somatic mutations are abundant, it is computationally hard to efficiently prioritize the identified neoantigen candidates according to their ability of activating the T cell immuno-response and numerous prioritization or prediction approaches have been proposed to address this issue. However, none of the existing approaches considers the original DNA loci of the neoantigens in the 3D genome perspective, to the best of our knowledge. In this work, we retrospect the DNA origin of the immune-positive and immune-negative neoantigens in the context of 3D genome and discovered that 1) Immuno-positive neoantigens’ corresponding DNA tend to cluster in some chromosomes (intra-chromosome) and tend to cluster genome-wise (inter-chromosome); 2) Immuno-active neoantigens’ corresponding DNA tend to belong to active chromosomal compartments (compartment A) in some chromosomes; 3) Immuno-active neoantigens’ corresponding DNA tend to locate at specific regions in the 3D genome. We believe that the 3D genome information, combining advanced machine learning[27-29] and feature selection technologies[30-32], will help more precise neoantigen prioritization and discovery, and will eventually benefit precision medicine in cancer immunotherapy.

## References

[1] T. N. Schumacher and R. D. Schreiber, “Neoantigens in cancer immunotherapy,” Science, vol. 348, no. 6230, pp. 69–74, Apr 3 2015.

[2] M. Yarchoan, B. A. Johnson, 3rd, E. R. Lutz, D. A. Laheru, and E. M. Jaffee, “Targeting neoantigens to augment antitumour immunity,” Nat Rev Cancer, vol. 17, no. 4, pp. 209–222, Apr 2017.

[3] S. G. O’Brien et al., “Imatinib compared with interferon and low-dose cytarabine for newly diagnosed chronic-phase chronic myeloid leukemia,” N Engl J Med, vol. 348, no. 11, pp. 994–1004, Mar 13 2003.

[4] H. Hackl, P. Charoentong, F. Finotello, and Z. Trajanoski, “Computational genomics tools for dissecting tumour-immune cell interactions,” Nat Rev Genet, vol. 17, no. 8, pp. 441–58, Jul 4 2016.

[5] C. Lundegaard, K. Lamberth, M. Harndahl, S. Buus, O. Lund, and M. Nielsen, “NetMHC-3.0: accurate web accessible predictions of human, mouse and monkey MHC class I affinities for peptides of length 8-11,” Nucleic Acids Res, vol. 36, no. Web Server issue, pp. W509–12, Jul 1 2008.

[6] M. Nielsen and M. Andreatta, “NetMHCpan-3.0; improved prediction of binding to MHC class I molecules integrating information from multiple receptor and peptide length datasets,” Genome Med, vol. 8, no. 1, p. 33, Mar 30 2016.

[7] V. Jurtz, S. Paul, M. Andreatta, P. Marcatili, B. Peters, and M. Nielsen, “NetMHCpan-4.0: Improved Peptide-MHC Class I Interaction Predictions Integrating Eluted Ligand and Peptide Binding Affinity Data,” J Immunol, vol. 199, no. 9, pp. 3360–3368, Nov 1 2017.

[8] S. J. Blevins et al., “How structural adaptability exists alongside HLA-A2 bias in the human alpha beta TCR repertoire,” Proceedings of the National Academy of Sciences of the United States of America, vol. 113, no. 9, pp. E1276–E1285, Mar 1 2016.

[9] T. P. Riley et al., “T cell receptor cross-reactivity expanded by dramatic peptide-MHC adaptability,” Nature Chemical Biology, vol. 14, no. 10, pp. 934–+, Oct 2018.

[10] Y. Wang et al., “How an alloreactive T-cell receptor achieves peptide and MHC specificity,” Proceedings of the National Academy of Sciences of the United States of America, vol. 114, no. 24, pp. E4792–E4801, Jun 13 2017.

[11] G. L. Zhang, H. H. Lin, D. B. Keskin, E. L. Reinherz, and V. Brusic, “Dana-Farber repository for machine learning in immunology,” J Immunol Methods, vol. 374, no. 1–2, pp. 18–25, Nov 30 2011.

[12] R. Vita et al., “The Immune Epitope Database (IEDB): 2018 update,” Nucleic Acids Res, vol. 47, no. D1, pp. D339–D343, Jan 8 2019.

[13] S. K. Gupta, T. Jaitly, U. Schmitz, G. Schuler, O. Wolkenhauer, and J. Vera, “Personalized cancer immunotherapy using Systems Medicine approaches,” Briefings in Bioinformatics, vol. 17, no. 3, pp. 453–467, May 2016.

[14] K. C. Parker, M. A. Bednarek, and J. E. Coligan, “Scheme for ranking potential HLA-A2 binding peptides based on independent binding of individual peptide side-chains,” J Immunol, vol. 152, no. 1, pp. 163–75, Jan 1 1994.

[15] M. M. Schuler, M. D. Nastke, and S. Stevanovikc, “SYFPEITHI: database for searching and T-cell epitope prediction,” Methods Mol Biol, vol. 409, pp. 75–93, 2007.

[16] M. Moutaftsi et al., “A consensus epitope prediction approach identifies the breadth of murine TCD8+-cell responses to vaccinia virus,” Nature Biotechnology, vol. 24, no. 7, pp. 817–819, Jul 2006.

[17] E. Karosiene, C. Lundegaard, O. Lund, and M. Nielsen, “NetMHCcons: a consensus method for the major histocompatibility complex class I predictions,” Immunogenetics, vol. 64, no. 3, pp. 177–86, Mar 2012.

[18] E. Karosiene, M. Rasmussen, T. Blicher, O. Lund, S. Buus, and M. Nielsen, “NetMHCIIpan-3.0, a common pan-specific MHC class II prediction method including all three human MHC class II isotypes, HLA-DR, HLA-DP and HLA-DQ,” Immunogenetics, vol. 65, no. 10, pp. 711–24, Oct 2013.

[19] T. Trolle et al., “Automated benchmarking of peptide-MHC class I binding predictions,” Bioinformatics, vol. 31, no. 13, pp. 2174–81, Jul 1 2015.

[20] B. Peters et al., “A community resource benchmarking predictions of peptide binding to MHC-I molecules,” PLoS Comput Biol, vol. 2, no. 6, p. e65, Jun 9 2006.

[21] P. Wang, J. Sidney, C. Dow, B. Mothe, A. Sette, and B. Peters, “A systematic assessment of MHC class II peptide binding predictions and evaluation of a consensus approach,” PLoS Comput Biol, vol. 4, no. 4, p. e1000048, Apr 4 2008.

[22] Y. Shi, X. B. Su, K. Y. He, B. H. Wu, B. Y. Zhang, and Z. G. Han, “Chromatin accessibility contributes to simultaneous mutations of cancer genes,” Sci Rep, vol. 6, p. 35270, Oct 20 2016.

[23] J. R. Dixon et al., “Topological domains in mammalian genomes identified by analysis of chromatin interactions,” Nature, vol. 485, no. 7398, pp. 376–80, Apr 11 2012.

[24] G. M. Boratyn et al., “BLAST: a more efficient report with usability improvements,” Nucleic Acids Res, vol. 41, no. Web Server issue, pp. W29–33, Jul 2013.

[25] D. W. Huang, B. T. Sherman, and R. A. Lempicki, “Systematic and integrative analysis of large gene lists using DAVID bioinformatics resources,” Nature Protocols, vol. 4, no. 1, pp. 44–57, 2009.

[26] W. J. Xie, L. Meng, S. Liu, L. Zhang, X. Cai, and Y. Q. Gao, “Structural Modeling of Chromatin Integrates Genome Features and Reveals Chromosome Folding Principle,” Sci Rep, vol. 7, no. 1, p. 2818, Jun 6 2017.

[27] V. Mnih et al., “Human-level control through deep reinforcement learning,” Nature, vol. 518, no. 7540, pp. 529–33, Feb 26 2015.

[28] Y. Yuan et al., “DeepGene: an advanced cancer type classifier based on deep learning and somatic point mutations,” BMC Bioinformatics, vol. 17, no. Suppl 17, p. 476, Dec 23 2016.

[29] Y. Yuan et al., “Cancer type prediction based on copy number aberration and chromatin 3D structure with convolutional neural networks,” BMC Genomics, vol. 19, no. Suppl 6, p. 565, Aug 13 2018.

[30] Z. P. Cai and L. Z. Xu, “Using gene clustering to identify discriminatory genes with higher classification accuracy,” Bibe 2006: Sixth Ieee Symposium on Bioinformatics and Bioengineering, Proceedings, pp. 235–+, 2006.

[31] K. Yang, Z. Cai, J. Li, and G. Lin, “A stable gene selection in microarray data analysis,” BMC Bioinformatics, vol. 7, p. 228, Apr 27 2006.

[32] Z. Cai, T. Zhang, and X. F. Wan, “A computational framework for influenza antigenic cartography,” PLoS Comput Biol, vol. 6, no. 10, p. e1000949, Oct 7 2010.

